# Succession comprises a sequence of threshold-induced community assembly processes towards multidiversity

**DOI:** 10.1101/2021.05.18.444608

**Authors:** Maximilian Hanusch, Xie He, Victoria Ruiz-Hernández, Robert R. Junker

**Affiliations:** Department of Biosciences, Paris Lodron University Salzburg, 5020 Salzburg, Austria; Evolutionary Ecology of Plants, Department of Biology, Philipps-University Marburg, 35043 Marburg, Germany

**Keywords:** succession, community assembly, ecological threshold, multidiversity, regime-shift, multi-betadiversity, biotic interactions

## Abstract

Research on successions and community assembly both address the same processes such as dispersal, species sorting, and biotic interactions but lack unifying concepts. Recent theoretical advances integrated both research lines proposing a sequence of stochastic and niche-based processes along successional gradients. Shifts in assembly processes are predicted to occur abruptly once abiotic and biotic factors dominate over dispersal as main driver. Considering the multidiversity composed of five organismal groups including plants, animals, and microbes, we empirically show that stochastic dispersal-dominated community assembly is replaced by environmental filters and biotic interactions after around 60 years of succession in a glacier forefield. The niche-based character of later successional processes is further supported by a decline in multi-beta-diversity. Our results may update concepts of community assembly by considering multiple taxa, help to bridge the gap between research on successions and community assembly, and provide new insights into the emergence of multidiverse and complex ecosystems.

## Introduction

Succession and community assembly research are neighboring fields in ecology, each with a long history of constructive debates and a large body of literature. Both research lines are founded on overlapping concepts considering processes such as dispersal, environmental filtering, biotic interactions, and stochasticity as structuring elements of local communities and ecosystems. Nevertheless, succession and community assembly research focus on different spatial and temporal scales, which may have hindered a mutual exchange of ideas (Chang and HilleRisLambers, 2016). Succession research relates to the initial development of ecosystems and communities over time. Community assembly studies, on the other hand, try to elucidate the past drivers of local and recent diversity patterns regardless of temporal components. Attempts to synthesize the two fields led to the development of an integrated conceptual framework of succession and community assembly dynamics. Here, drivers of community structure are considered from an explicitly temporal perspective, assuming a temporal sequence of assembly processes shifting from dispersal to niche-based and interaction-mediated processes along the course of succession (Chang and HilleRisLambers, 2016). The proposed framework also emphasizes the importance of threshold dynamics, predicting that gradual changes in the abiotic and biotic environment are followed by rapid shifts in the predominant processes that determine community assembly (Suding and Hobbs, 2009; Kadowaki *et al*., 2018). These shifts are expected to be associated with various community level responses such as changes in diversity, species composition and functionality (Scheffer *et al*., 2001; Suding and Hobbs, 2009; Chang and HilleRisLambers, 2016). Recent research suggests that sudden ecosystem shifts are not restricted to changes in species composition of single organismal groups but that local diversity patterns emerge from on-site interactions between multiple taxa (Lu and Hedin, 2019; Berdugo *et al*., 2020). Multidiversity, an aggregate measure of biodiversity that integrates the standardized diversities of multiple taxa (Allan *et al*., 2014), has been shown to reflect ecosystem functionality (Delgado-Baquerizo *et al*., 2020), species composition (Felipe-Lucia *et al*., 2020), and multitrophic interactions (Manning *et al*., 2015) more accurately than diversity measures considering only one taxon. Additionally, the responses of different organismal groups to successional gradients may differ either due to stochastic or deterministic drivers (Kaufmann, 2001; Blaalid *et al*., 2012; Fenton and Bergeron, 2013; Dini-Andreote *et al*., 2015; Måren *et al*., 2018; Junker *et al*., 2021) and thus conclusions about the primary assembly processes may be biased in studies considering a single or few taxa only (Pulsford, Lindenmayer and Driscoll, 2016).

The assumed changes of assembly processes along successions, a decrease in the importance of dispersal and a simultaneous increase in niche-based and interaction-based processes, have consequences for the taxonomic, functional and phylogentic composition of communities and thus beta diversity (Odum, 1969; Chang and HilleRisLambers, 2016). The framework by Chang & HilleRisLambers (2016) predicts an increase in functional and phylogentic diversity of local single-taxon communities in later successions due to niche differentiation and intensified biotic interactions. In a multidiversity context where multiple taxa are considered, increased interactions and thus potentially strong dependencies between taxa may result in aggregated co-occurrence patterns (Ohlmann *et al*., 2018), which may have consequences for the beta-diversity between local communities – and not necessarily for the functional and phylogenetic diversity within a community. For instance, interactions between plants and soil inhabiting microorganisms intensify with successional age (Chen *et al*., 2019) and regulate plant-animal interactions (Howard, Kao-Kniffin and Kessler, 2020), which is reflected in the co-occurence of the partners involved in these interactions. Such interactions may thus result in reduced species turnover across local assemblages in older successional stages. Yet, it is an ongoing debate whether ecological communities generally converge towards a core set of species over the course of succession (Lepš *et al*., 1991; Fukami *et al*., 2005) and studies on successional convergence have mostly been focused on singular taxa neglecting potential interactions and co-occurrences across organismal groups (Brown and Jumpponen, 2014; Castle *et al*., 2016; Chang *et al*., 2019). To specifically test the effect of interactions across taxa on species turnover, we introduce multi-betadiversity, an aggregated measure of taxonomic community dissimilarity. This index allows to test whether the expected increase in biotic interactions is reflected in a reduced compositional variation (i.e., reduced multi-betadiversity regarding several taxa) of the multidiverse community once niche-based and interaction-mediated assembly processes dominate over dispersal as main mechanisms shaping multidiversity.

Assuming that multidiversity more precisely reflects ecosystem responses and functions, we tested predictions deduced from contemporary succession hypotheses using a dataset on multidiversity comprising inventories of plants, animals, and microorganisms along a successional gradient. Most studies on multidiversity have been undertaken in well-developed ecosystems that have undergone severe anthropogenic alterations in the past and focused on the drivers of biodiversity decline (Allan *et al*., 2014; Manning *et al*., 2015; Felipe-Lucia *et al*., 2020). The mechanisms behind the successional emergence of multidiversity and ecosystem complexity under natural conditions, however, are yet poorly understood. Thus, considering multidiversity within a successional framework will help to gain new insights into the temporal dynamics of the processes that shape the initial emergence of biodiversity in natural ecosystems that comprise multiple organismal groups with unique and complementary ecological functions. We adopted the concept of multidiversity to empirically test predictions of the integrated framework of succession and community assembly dynamics on an ecological gradient of primary succession following glacial retreat in the Austrian Alps (Junker *et al*., 2020). We assessed the multidiversity and multi-betadiversity of five organismal groups, including vascular plants, bryophytes, invertebrates, and soil inhabiting fungi and bacteria on 110 plots spanning 170 years of primary succession. We estimated multidiversity by the sum of ranked normalized Shannon-diversities and calculated multi-betadiversity as the mean Bray-Curtis dissimilarities between plots regarding the composition of individual taxa (see methods). According to the conceptual framework by Chang & HilleRisLambers (2016), we predicted that community assembly will undergo a rapid shift that is induced by a sudden change of the biotic and abiotic environment along the successional gradient. We expect this shift to be associated with increased biotic interactions that result in a reduced compositional variation between plots of the multidiverse community.

Accordingly, we identified a threshold in multidiversity development (i.e., a shift from an increase of multidiversity with time to a rather stationary phase) and estimated the relative importance of dispersal, environmental and biotic drivers on the assembly of the multidiverse community before and after the threshold using path analysis. We further highlighted the more structured character of community assembly after the shift by comparing multi-betadiversity estimates of the early and late successional communities.

## Results

### Breaking point in multidiversity

Multidiversity follows a non-linear trajectory along the successional gradient. We screened the gradient for the predicted shift from a developing (characterized by an increase in multidiversity over time) to a more stationary ecosystem (characterized by non-monotonic variation in multidiversity over time) using breaking point analysis, which detects changes in the slope of associations. According to Groffman (2006) breaking points represent ecological thresholds, which is characterized by a rapid change of the slope in the association between time since deglaciation as explanatory variable and multidiversity as dependent variable in our study. We found such a breaking point by cross-validating a broken-stick model to alternative models that either did not include a breaking point or included a disjunct breaking point using Bayesian inference (Lindeløv, 2020). We then used Bayes Factors analysis to validate the exact location of the breaking point at plot number 44 which reflects about 60 years of succession (see Methods, Fig. 1, Supplementary Table 1). Resulting piecewise linear models showed a significant linear increase in multidiversity during the early successional stage and a stationary multidiversity (non-significant relationship with time) for the late successional stage (early: *t*_42_ = 5.77, *p* < 0.001, *r*^*2*^ = 0.44; late: *t*_64_ = -0.55, *p* = 0.58, *r*^*2*^ = 0.004; Fig.1). To assess whether the change in assembly processes is also reflected in the composition of the multidiverse community, we calculated the relative abundance of each taxon (see Methods) along the successional gradient and visualized it in a temporally ordered multi-taxa community table of species occurrences (Fig. 2). The threshold became also evident in the change of community composition with different sets of taxa before and after the breaking point. Prior to the threshold, communities mainly consist of species that reach their abundance optima early in succession. These pioneering species were soon accompanied by taxa with no clear preference of early or late successional stages leading to an increase in multidiversity over time. After the threshold, specialists for early successions are replaced by specialists for late successional stages, consequently multidiversity remained stationary over time (Fig. 2).

**Figure 1:**
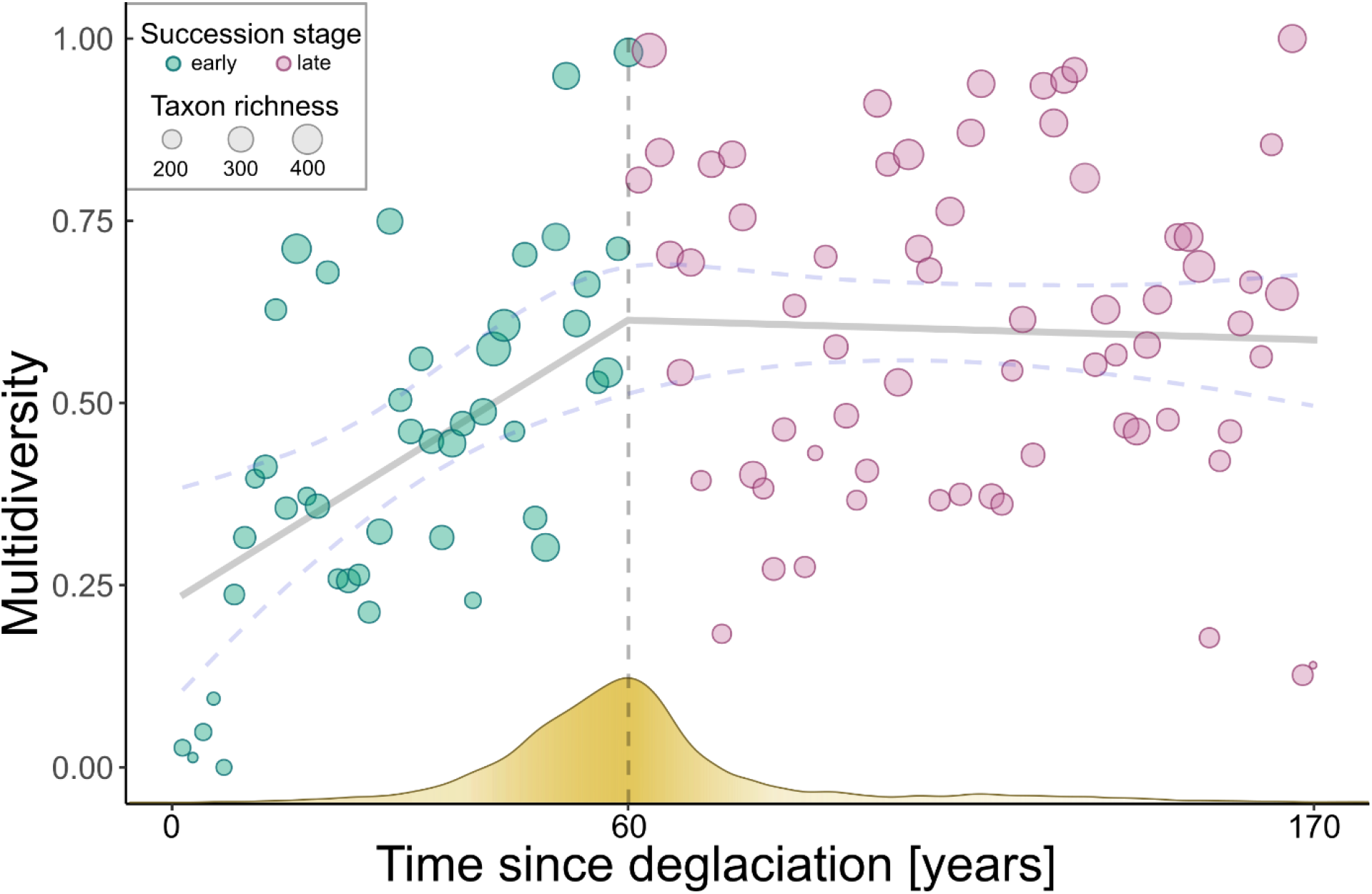
Multidiversity along the successional gradient. Multidiversity increases during the early successional stage and remains stationary as succession continues. The vertical dashed line indicates the estimated threshold in multidiversity development, which is the threshold in community assembly as predicted by the framework of Chang & HilleRisLambers (2016). The grey regression line indicates the linear correlation between multidiversity and time since deglaciation using a broken-stick-model. Blue dashed lines indicate the 2.5% and 97.5% quantiles of fitted values. The yellow curve on the x-axis resembles the posterior distribution for the estimate of the breaking point i.e., higher density of the frequency indicates a higher probability for the breaking point to be located at a given value.

**Figure 2:**
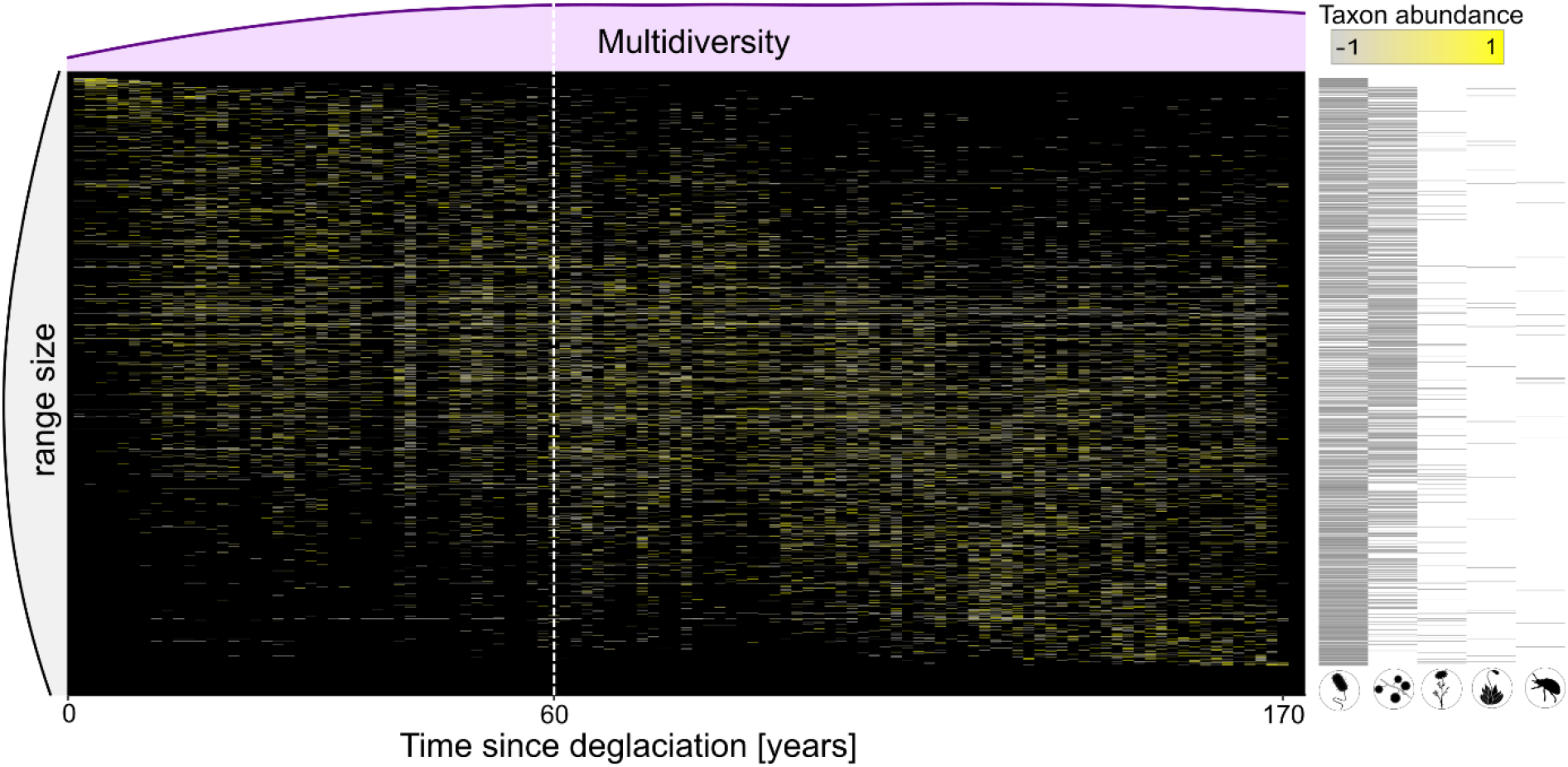
Single taxa occurrence along the successional gradient for each taxon that occurs on at least three plots along the gradient. Tiles indicate the relative abundance of each taxon at a given plot relative to the mean abundance of the taxon along the gradient. Taxa are ordered by the weighted average plot of their occurrence, i.e. taxa that reach their abundance optima early in succession appear in the top left and late successional specialist in the bottom right. Relative taxon abundance is color-coded and reflects z-scores with the mean abundance as 0 and one standard deviation as +1 or -1, rescaled between -1 and 1. Multidiversity is indicated as light pink density plot on top of the community composition plot. Grey density plot on the left side indicates the range size of the taxa, i.e whether their occurrence is restricted to a narrow range of the successional gradient or whether taxa occur along the whole gradient. Black bars on the right represent the organismal group of the taxa in each row, symbols of organismal groups are ordered by decreasing frequency; white dashed horizontal line marks the threshold.

### Drivers of multidiversity

To quantify the relative importance of dispersal and environmental drivers, we specified a path model in which time since deglaciation (as a proxy for dispersal events that occur at a random frequency as main driver (Lowe and McPeek, 2014)) and mean seasonal soil temperature (as an environmental factor that is important for diversity of several trophic levels (Ohler, Lechleitner and Junker, 2020)) act as exogenous variables and directly affect diversities of single organismal groups. The total effects of the dispersal and environmental drivers on multidiversity were estimated as indirect effects mediated through the organismal groups. We also tested the effects of other environmental variables, such as soil nutrient content and soil-pH value but could not detect any significant effects and thus removed these variables from the final model (see Methods and Supplementary tables 2-4). Biotic interactions were modeled as the residual covariance among the diversity values of all organismal groups. Community covariances can be interpreted as biotic interactions and especially positive covariances are indicative of facilitative effects within the community (Houlahan *et al*., 2008; Ranta *et al*., 2008). The path analysis provided good model fit (Model fit: p_chi-square_ > 0.05; CFI >0.95; TLI ≥ 0.9; SRMR < 0.09; RMSEA_early_ < 0.05; RMSEA_late_ = 0.066 with lower bound of confidence interval = 0) and revealed different influences of the exogenous variables between the early and late successional environments (Figure 3, Supplementary Table 5). During early succession, time since deglaciation had a strong positive effect on the diversity of bryophytes (β = 0.65, p < 0.01), arthropods (β = 0.61, p < 0.01), and bacteria (β = 0.31, p = 0.04), whereas there was no significant effect of temperature on any organismal group. Time since deglaciation also had a significant positive indirect effect on total multidiversity (β = 0.74, p < 0.01) during early succession, whereas temperature did not affect multidiversity (β = 0.01, p = 0.93, Figure 3a). Furthermore, diversities of the five organismal groups varied independently of each other suggesting little or no mutual influences. In late succession, mean soil temperature had a significant positive direct effect on the diversity of bryophytes (β = 0.30, p = 0.01), yet there was a significantly positive indirect effect on multidiversity mediated through all organismal groups (β = 0.29, p = 0.01). Plot age had no longer an effect on multidiversity (β = -0.01, p = 0.89) and we detected positive residual covariances between vascular plant (β = 0.26, p = 0.04), invertebrate (β = 0.25, p = 0.05) and bacterial (β = 0.28, p = 0.02) diversities with fungal diversity, suggesting mutual influences between taxa (Figure 3b).

**Figure 3:**
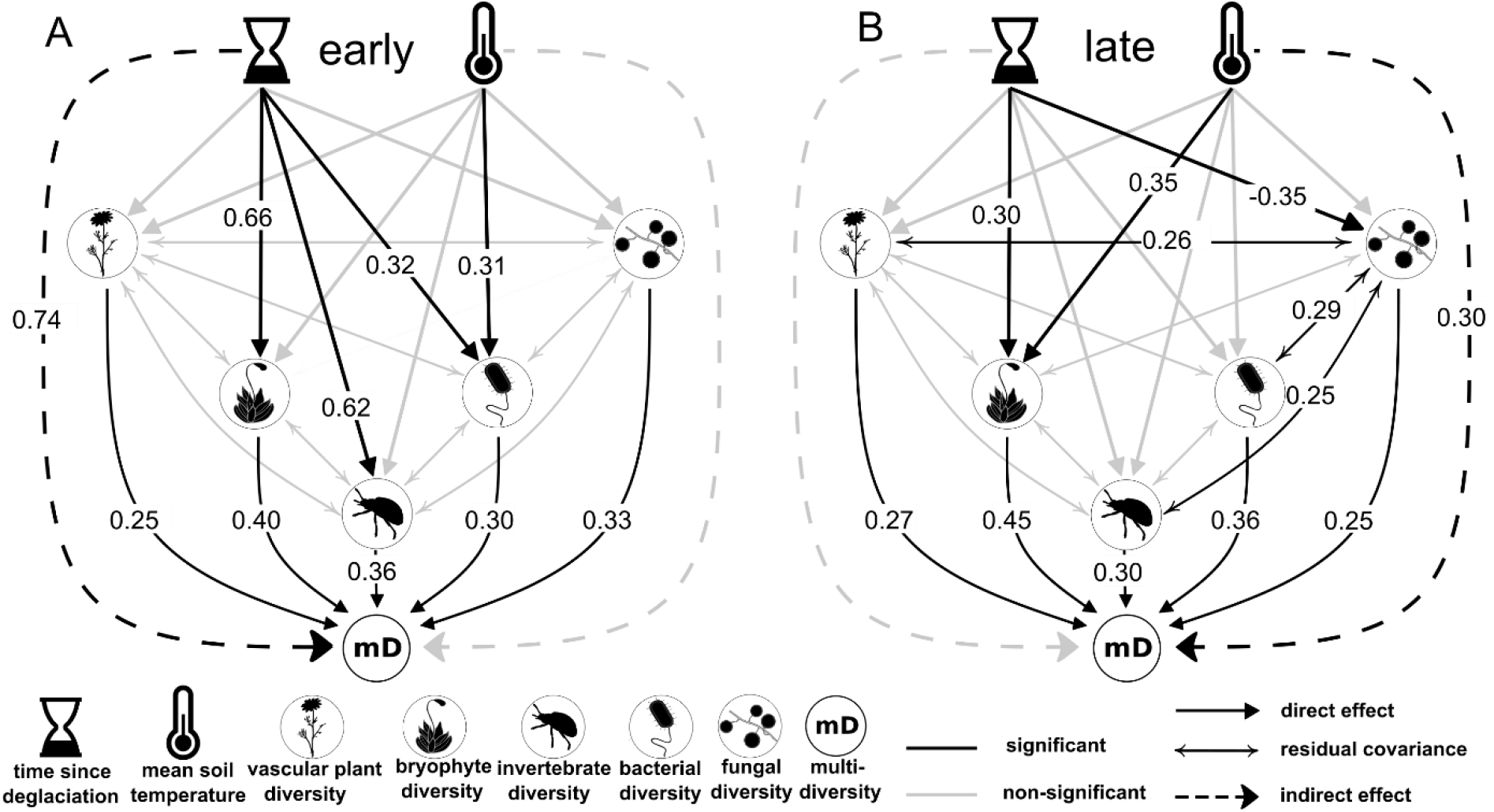
Path analysis regarding the effects of stochastic and deterministic drivers on the multidiverse community for the early and late successional stages. Time since deglaciation and mean seasonal temperature act as exogenous variables with directed effects on the diversities of vascular plants, bryophyte diversity, invertebrates, and soil inhabiting fungi and bacteria. Indirect effects of exogenous variables on multidiversity were calculated by mediation analysis through the direct effects of the organismal groups on multidiversity. Biotic interactions are modeled as covariances between organismal groups. Numbers represent standardized path coefficients and are given for significant paths only.

### Multibetadiversity

We predicted that the shift from dispersal dominated stochastic events in early successional stages to niche-based processes in late successional stages is associated with lower beta-diversity among communities after the threshold. To test this prediction, we calculated a measure of total community dissimilarity, multi-betadiversity (*mbD*) for both successional stages by bootstrapping Bray-Curtis-similarities of the multidiverse community (see Methods). Despite different trajectories of betadiversity for different organismal groups -an increase in dissimilarity for bryophyte, and a decrease for those groups that show positive community covariances in the path model in the old successional stage (bacteria, fungi, and vascular plant), and no detectable change for invertebrate communities - multibetadiversity was significantly lower in the late successional stage (mean ± SD; *mbD*_early_ = 0.73 ± 0.14; *mbD*_late_ = 0.71 ± 0.13; Fig. 4).

**Figure 4:**
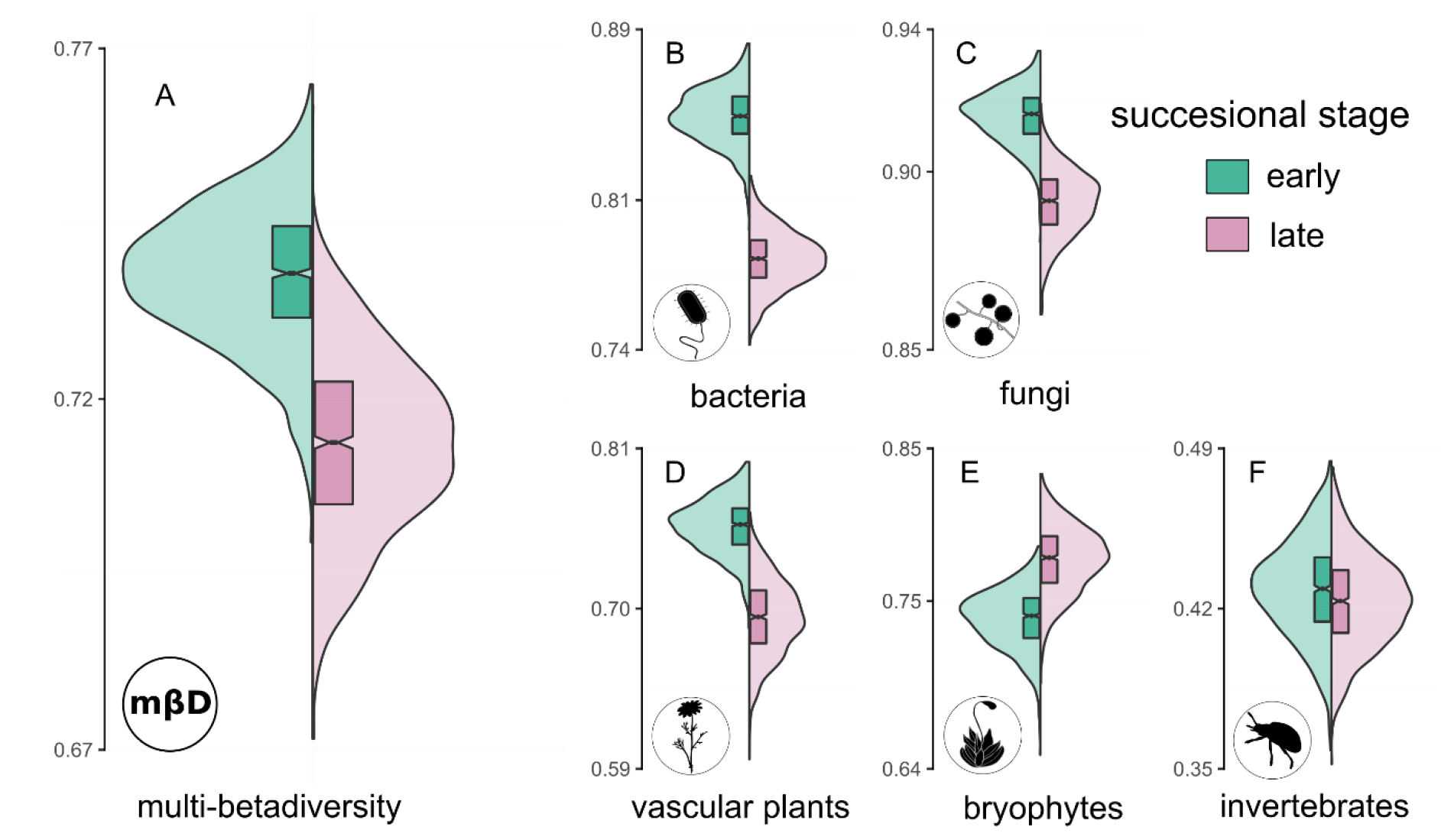
Betadiversity values for each organismal group and community-wide multi-betadiversity values for the early and late successional stages. Mean Bray-Curtis dissimilarities were used to calculate betadiversity and multi-betadiversity values. Green violin plots resemble the dissimilarity values of the early successional stage (n=44 plots), rose violin plots indicate the late successional stage (n=66 plots). Violin plots indicate the distribution of the mean dissimilarities for n=1000 bootstrap draws. Boxplots indicate the upper and lower quartile of the estimates, and the size of the notch represents the confidence interval of the median. A) mean of all organismal groups, i.e., multi-betadiversity B) bacteria C) fungi D) vascular plants E) bryopyhtes F) invertebrates

## Discussion

Our study is an empirical test of the conceptual framework integrating succession and community assembly dynamics proposed by Chang & HilleRisLambers (2016)^1^ and modified here in the context of multidiverse communities. We showed that threshold dynamics play an important role in the assembly of multidiverse communities under natural conditions and that these thresholds are associated with substantial changes in the community assembly processes. So far, the importance of threshold dynamics has been recognized for anthropogenically altered ecosystems either as catastrophic shifts with respect to climate change (Scheffer *et al*., 2001; Berdugo *et al*., 2020) or for restoration efforts of anthropogenically altered landscapes (Suding and Hobbs, 2009; Kadowaki *et al*., 2018). We revealed a threshold after about 60 years of ecosystem development when multidiversity became stationary. Significant thresholds that are marked by a steep increase followed by stationarity after 40 to 60 years of succession appear to be a generalizable pattern that occurs in various aspects of ecosystem development, such as plant, invertebrate, and microbial diversities and functionality in glacier forefields in Europe and Northern America (Kaufmann, 2001; Tscherko *et al*., 2003; Raffl *et al*., 2006; Schütte *et al*., 2010; Dong *et al*., 2016). Thus, acknowledging the effects of threshold dynamics during the development of natural multidiverse ecosystems, and their universality in ecological systems provides valuable insights into the patterns and processes of initial ecosystem development.

Our results indicate a strongly dispersal driven assembly in the first 60 years of succession and a pronounced influence of the environment as well as increased biotic interactions afterwards. Although different organismal groups follow individual assembly trajectories during primary succession, multidiversity is mainly promoted by stochastic drivers during the initial phase of ecosystem development, most assumably by heterogenous dispersal events. Over the course of succession, stochasticity is replaced by environmental filtering and biotic interactions as the structuring mechanism of multidiversity. Our data do not allow to directly test for mutual influences between organismal groups on the taxon level, but we find a strong pattern of positive community covariances after accounting for environmental variation that is indicative of facilitative processes and biotic dependencies within the community (Houlahan *et al*., 2007; Ranta *et al*., 2008). A previous study in the same glacier forefield showed that after more than 50 years of ecosystem development, the soil microbial community is mainly supported by carbon from recent plant production, whereas during the initial stage of ecosystem development, the microbial community is mainly sustained by ancient carbon sources (Bardgett *et al*., 2007). The time point of the shift in the main carbon source largely coincides with our delineated threshold in multidiversity development. The increased interdependence of the soil microbial and plant community during the later successional stage is also reflected in the positive covariances of vascular plant, fungal and bacterial diversities in our path model. Although bryophytes have been shown to closely interact with epiphytic microorganisms after glacial retreat (Arróniz-Crespo *et al*., 2014), potential interactions of bryophytes with microorganisms may not be detectable in our path-model as we sampled soil microbial communities and not all substrate types colonized by bryophytes in our study site (e.g. rocks, scree and litter).

The threshold-mediated character of successional processes is further supported by a pronounced decline in multibeta-diversity that is indicative of an increased dependence structure among the organismal groups. In line with that assumption, we could only detect a lower beta-diversity in those organismal groups that also show positive community covariances in the path model. Although co-occurrence patterns must not necessarily reflect biotic interactions (Blanchet, Cazelles and Gravel, 2020), this finding strengthens the assumption of biotic interactions as an underlying cause for a reduced beta-diversity between plots in later successional stages, which finds support in studies on plant community assembly (Fukami and Nakajima, 2013; Martínez-García *et al*., 2015). The literature suggests the increasing importance of interactions in community development trajectories. One explanation put forward is that with increasing niche differentiation, energy and nutrient flows in an ecosystem follow more complex biochemical pathways that necessitate biotic interactions to retain nutrients in the ecosystem through closed mineral cycles (Odum, 1969; Chang and HilleRisLambers, 2016). Our results support the increasing importance of biotic interactions as in the late successional stage, our path model revealed positive community covariances between the soil microbial, vascular plant, and arthropod communities that may represent such complex nutritional cycles - comprising primary producers, heterotrophic organisms, and decomposers - that are limited by energy influx (reflected as soil temperature in our model). For the organismal groups forming these putative cycles, we find a reduced betadiversity between the plots that indicates a strong interrelatedness of individual taxa after the threshold, which may lead to re-occurring core assemblages of species of several taxa. These assumptions and the threshold suggested by our data after 60 years are further supported by the study of Bardgett et. al (2007) that revealed a high dependence of heterotrophic microorganisms on plant communities in the Ödenwinkel forefield after a similar number of years. We thus recommend extending the integrative framework of succession and community assembly for multiple interacting taxa that mutually shape their diversity and composition and thus cause a reduction of betadiversity.

Understanding the relative importance and temporal dynamics of deterministic and stochastic processes is a key challenge in community ecology, especially in natural systems and has been in the center of broad debates among ecologists. Using the multidiversity and multi-betadiversity approach allows us to comprehensively understand the processes leading to stable, resilient, and complex ecosystems, which may remain vague in single-taxon approaches. Thus, our study contributes to a synthesis of community ecological theories into succession research, acknowledging the fundamental importance of abrupt state shifts in natural ecosystems. These results are not only a proof of concept but also emphasize that succession is a multi-faceted rather than a linear process that comprises a sequence of assembly processes towards multidiversity and ecosystem complexity.

## Methods

### Study Design

The study was conducted in the long-term ecological research platform Ödenwinkel which was established in 2019 in the Hohe Tauern National Park, Austria (Dynamic Ecological Information Management System – site and dataset registry: https://deims.org/activity/fefd07db-2f16-46eb-8883-f10fbc9d13a3, last access: March 2021). A total of *n* = 135 permanent plots was established within the glacier forefield of the Ödenwinkelkees which was covered by ice at the latest glacial maximum in the Little Ice Age around 1850. For this study, we used a subset of *n* = 110 plots with complete datasets with all biotic and abiotic variables available. The plots represent a successional gradient spanning over 1.7km in length and were evenly distributed within the glacier forefield at an altitude ranging from 2070 to 2170 m a.s.l. Plots were defined as squares with an area of 1 m^2^ and were all oriented in the same cardinal direction. Further details on the design of the research platform, exact plot positions and details on the surrounding environment, as well as on the historical glacial extent can be found in Junker et al. (2020).

### Sampling

In 2019, we identified all vascular plant and bryophyte species present on the plots and estimated their cover with a resolution of 0.1 %. We sampled above-ground arthropod diversity by installing two pitfall traps on each plot. Traps were set active for a total of *n* = 7 days. The abundance of all arthropods, excluding Collembola and Acari, larger than 3 mm was counted. The abundance of Collembola and Acari and of animals smaller than 3mm was estimated based on random samples of aliquots of the total sample. All arthropods and other animals are identified to the order level. Soil inhabiting bacteria and fungi were sampled from soil cores from an approximate depth of 3 cm, as soil development in the proglacial study area was limited to the top layers of the pedosphere. Soil-microbiome samples were analyzed by next-generation sequencing and microbiome profiling of isolated DNA was performed by Eurofins Genomics (Ebersberg, Germany). Prior to the statistical analysis of microbial communities, we performed a cumulative sum scaling (CSS) normalization (R package “metagenomeSeq” v1.28.2, Paulson *et al*. 2013) on the count data to account for differences in sequencing depth among samples. Detailed information on the sampling strategies of all organismal groups can be found in Junker et al. (2020). Temperature measurements were done by installing temperature loggers (MF1921G iButton, Fuchs Elektronik, Weinheim, Germany) 10 cm north of each plot centre, at the same depth of 3 cm at which the microbial samples were taken. Mean temperature of the growing season was calculated based on the recordings ranging from 26th of June to 16th of September representing the period in which the plots were free of permanent snow cover before and after the winter 2019/2020. In 2020, soil samples were taken and soil nutrients (Ca, P, K, Mg and total N_2_) as well as soil pH were measured on all plots by AGROLAB Agrar und Umwelt GmbH (Sarstedt, Germany). Soil nutrient analysis was performed according to the Ö-Norm Boden: L 1087: 2012-12 (K and P - mg/1000g), L 1093: 2010-12 (Mg - mg/1000g) and L 1083: 2006-04 (pH). Total N2 (%) was determined according to the DIN EN 16168: 2012-11. Detailed information on the sampling strategy can be found in Junker et. al (2020).

### Calculation of multidiversity

Multidiversity is defined as the cumulative diversity of a number of taxonomic groups (Allan *et al*., 2014). The multidiversity of the *n* = 110 plots composed of the diversities of vascular plants, bryophytes, invertebrates, fungi, and bacteria present in each plot and was defined as follows: First, we calculated the Shannon diversities (H) of each of the taxonomic groups in each plot. Second, we ranked the plots by increasing diversity for each of the five taxonomic groups individually, i.e., for each plot we received *n* = 5 ranks. The mean rank of each plot was defined as multidiversity *mD*. For better interpretability of the index, we then normalized the mean ranks as x’ = ((x - min(x)) / (max(x) - min(x)) to scale them between zero and one. Ranks were used to give the same weighting to each taxonomic group despite deviations in the absolute values of Shannon diversity and to reduce the impact of outliers in *mD* values. To account for differences in the sequencing depth of the microbial raw dataset, we performed multiple rarefactioning prior to the calculation of Shannon diversity by averaging the results of *n* = 999 iterations (R package “rtk” v0.2.5.7; Saary *et al*., 2017) instead of using the CSS-normalized dataset.

### Breaking point analysis

According to the hypothesis of community assembly dynamics (Chang & HilleRisLambers 2016) and by visual inspection of the relationship between multidiversity and successional age of the plots we expected two different stages of community establishment along the successional gradient. These stages are separatable by the transition from a developing (characterized by an increase in multidiversity over time) to a more stationary ecosystem (characterized by non-monotonic variation in multidiversity over time). Such non-linear associations are suggestive of regime-shifts of the ecosystem and can be interpreted as ecological thresholds (Groffman *et al*., 2006). Piecewise regression models have been shown to be the most suitable tool for the detection of ecological thresholds in natural systems, as they are able to correctly estimate the probability, as well as the number and position of ecological thresholds (Ficetola and Denoël, 2009). We screened the successional gradient for the existence of a threshold by comparing four different models with *mD* as the dependent and plot index number (surrogate for plot age) as the independent variable using the R-package “mcp” v0.3.0.9 (Lindeløv, 2020). The mcp-method fits piecewise regression models with a pre-defined number of breaking points and is based on Bayesian inference. We specified a base model m1 (*mD*-values remain constant along the gradient) and compared it to three alternative models (m2 = constant linear increase of *mD*-values without a breaking point, m3 = one breaking point in *mD*-values with a segregated slope (abrupt-threshold-model), m4 = one breaking point in *mD*-values with a joined slope (“broken-stick” or smooth-threshold-model). For each model, we separately ran three Markov Chain Monte Carlo estimators with a uniform prior for a total of n = 11000 generations while discarding the first n = 1000 generations as burn-in. Model convergence was estimated by visually inspecting the trace plots and checking that all model parameters reached stationarity. We then compared the predictive performance of the four models using leave-one-out-cross-validation and confirmed the exact position of the breaking point using Bayes Factors. Both validation methods can be applied for a robust and accurate testing of competing hypotheses in ecological datasets (Aho, Derryberry and Peterson, 2014; Conn *et al*., 2018; Hanusch *et al*., 2020).

### Community taxa occurrence

To visualize the distribution of individual taxa and thus the community wide taxonomic turnover along the successional gradient, we estimated the abundance optimum of each taxon that occurred on at least three plots. Abundance optima were estimated by calculating the mean plot of occurrence weighted by the abundance of each taxon (i.e., cover for vascular plants and bryophytes, individual count for invertebrates and CSS-normalized read number for microorganisms). We calculated the relative abundance per plot for each taxon by using z-scores of abundances. For each taxon, the mean abundance was set as 0 with one standard deviation as +1 or -1. Resulting z-scores were then rescaled between -1 and 1. Range size was estimated by calculating the variance of occurrence plots (i.e., the span of plots on which a taxon was found along the gradient) weighted by the abundance of the taxon on the respective plots.

### Path analysis

We used path-analysis i) to model the influence of the abiotic environment on the diversities of all taxonomic groups individually ii) to estimate the strength of community covariances between the diversities of those groups iii) to infer the effect sizes of the diversity of each group on total *mD* and iv) estimate the strength of indirect effects of the exogenous variables that are mediated through the organismal groups on *mD*. We built separate models with identical structure for the early (*n* = 44 plots) and late (*n* = 66 plots) successional stages. The stages were delineated by the threshold identified in the breakpoint analysis. Exogenous variables in the model were time since deglaciation and mean soil temperature of the growing season. Time since deglaciation reflects the plot age and can be seen as a proxy for the increasing chance of heterogenous dispersal events that occur over time. The mean temperature of the growing season is an estimate for environmental heterogeneity and has been shown to affect diversity directly and indirectly on various trophic levels (Ohler, Lechleitner and Junker, 2020). All variables were scaled by subtracting the mean and dividing through the standard deviation. We first ran a full model that included soil pH and soil nutrient content as additional exogenous variables. None of the two variables showed a significant direct effect on any organismal group or a significant indirect effect on multidiversity during either the young or late successional stage. Accordingly, we stepwise removed the two variables from the model while cross-checking whether the effect strength of one variable became significant in the absence of the other variable. As no significant effects occurred or the model fit significantly decreased, we decided to remove both variables from the final model. Within the final model, the strength of residual covariance between the diversities of all taxonomic groups was estimated while accounting for influences of time since deglaciation and temperature. If corrected for an underlying common cause, such as environmental autocorrelation, strong covariances between members of a community can be interpreted as biotic interactions and especially positive community covariances are indicative of facilitative effects within the community (Houlahan *et al*., 2008; Ranta *et al*., 2008). All path-models were estimated using the R-package “lavaan” v0.6-7 (Rosseel, 2012).

### Multibetadiversity

We defined Multibetadiversity (*mbD*) as the cumulative averaged pairwise-dissimilarity across a number of organismal groups. First, we split the total community composition data table (sites x species, *n* = 110 plots) for each organismal group into two tables containing the plots before (*n* = 44 plots) and after (*n* = 66 plots) the threshold. Second, we performed Bootstrapping (BS) on the subset tables for each of the *n* = 5 organismal groups individually. BS was done for each organismal group on both successional stages separately and was necessary to account for differences in sample sizes (i.e., number of plots) of the successional stages. Further, BS provides more robust inferential results and allows to obtain confidence intervals for the resulting beta diversity estimates. Each BS-replicate consisted of the following steps: For the early and late successional stage separately, we generated *n* = 1000 new community tables per organismal group by randomly sampling the communities of *n* = 44 plots (number of plots in the early successional stage) allowing replacement draws. We then calculated the mean of the pairwise Bray-Curtis dissimilarities for each of the bootstrapped community tables. Subsequently, we calculated *mbD* as the mean of the *n* = 5 dissimilarity values per BS-replicate.

## Contributions

R.R.J. conceived and initiated the study. M.H., X.H., V.R.H. and R.R.J designed the study and conducted field work. M.H. performed the processing and analysis of the data with main inputs from R.R.J. and X.H.. M.H. and R.R.J. drafted the initial version of the manuscript. All authors contributed critically to the interpretation of the results, revising, and approving the final version of the manuscript.

## Data availability

Raw sequences of next-generation 16S rRNA gene amplicon sequencing are available at the NCBI Sequence Read Archive (SRA) under the BioProject accession PRJNA701884 and PRJNA701890. Raw floristic and zoological data will be made available before publication.

## Code availability

R scripts will be made available before publication.

## Acknowledgements

We thank the Hohe Tauern National Park Salzburg administration and the Rudolfshütte for organizational and logistic support, the governing authority Land Salzburg for the permit to conduct our research (permit no. 20507-96/45/7-2019), Jan-Christoph Otto, Tobias Seifert, and Anna Vojtkó for help in the field. Hamed Azarbad, Lisa-Maria Ohler and Verena Zieschank provided valuable comments to improve the study. This research has been supported by the Austrian Science Fund (FWF), which provided funding to Robert R. Junker (grant no. Y1102).

## Corresponding author

Correspondence and requests for materials should be addressed to Robert R. Junker.

